## Supplementary information for "Modeling nearshore-offshore water exchange in Lake Ontario"

Electronic Supplementary Marterial:

The following five parts, listed below, provide supportive information cited in the manuscript.

- Part I. Model Setup,
- Part II. Calibration and validation
- Part III. Additional data
- Part IV. Bottom slope analysis
- Part V. Field Observations

Part I. Model setup

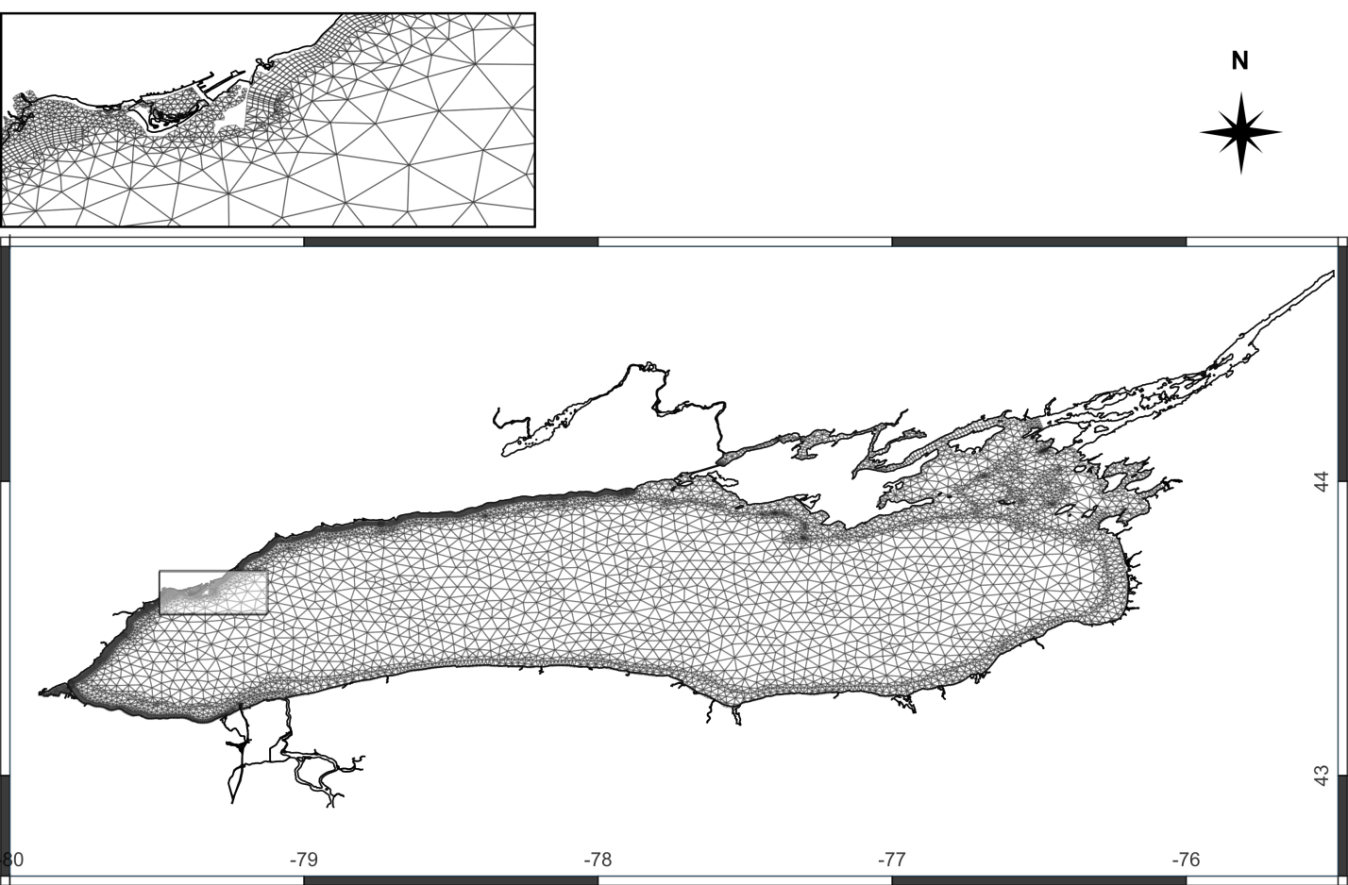

**S1 Fig.** Mike 3 flexible mesh for the entire Lake Ontario

Lake Ontario’s currents and thermal structure were simulated using DHI’s MIKE 3 (DHI, 2021), available at <https://www.mikepoweredbydhi.com>. MIKE 3’s hydrodynamic module is based on the numerical solution of the three-dimensional incompressible Reynolds averaged Navier-Stokes equations invoking the assumptions of Boussinesq and of hydrostatic pressure (DHI, 2021). MIKE 3 and its siblings MIKE 21 and MIKE Hydro (provided under the umbrella of MIKE Zero tools) are used in a wide range of applications (DHI, 2023). The modelling system is based on the conservation of mass and momentum in three dimensions of a Newtonian fluid. The flow is decomposed into mean quantities and turbulent fluctuations and the closure problem is solved through the Boussinesq eddy viscosity concept relating the Reynolds stresses to the mean velocity field, using a combination of the Smagorinsky formulation for the horizontal direction and a k-ε formulation for the vertical direction. The turbulence models are all solved in an explicit manner except for the vertical k-ε model, which is solved by an implicit scheme. To handle density variations, the equations for conservation of salinity and temperature are included. A flexible unstructured mesh was created with the Mesh Generator software provided in MIKE Zero. It contained over 15000 triangular elements in the horizontal plane, and 16 sigma layers up to the depth of 40 metres and an additional set of 14 z-level layers for deeper areas in the vertical direction. The bathymetry was prepared from a combination of data from measurements done by the Ontario Ministry of the Environment, Conservation and Parks (MECP) field crew and data available at the National Oceanographic and Atmospheric Administration (NOAA) website (<https://www.ngdc.noaa.gov/mgg/greatlakes/ontario.html>).

The heat exchange considered latent heat, sensible heat, short and long wave radiation, and atmospheric conditions, which were input directly from the NCAR data. Fluxes of latent and sensible heat were calculated in the model using the options available within the MIKE 3 modelling framework. The shortwave radiation was attenuated in the water column using the non-normalized Beer’s law corresponding to close to oligotrophic conditions (beta coefficient 0.25), which was assumed that reflects the condition in most of the lake (Weiskerger et al., 2018). The baroclinic equations time step was set to 30 seconds, which was considered adequate for numerical stability, and for the shallow water equations we used the low-order, fast algorithm options for both the time integration and space discretization. This compromise worked well to keep the simulation time around 7 days for a year of simulation, without noticing degradations in the quality of the results. The temperature equations and the Transport Module have used the same solution technique settings. ECCC and the NOAA deploy meteorological buoys that measure meteorological data and surface water parameters that include wave data. We used the wave data from ECCC buoy Stations C45135, C45139 and C45159 (<https://www.meds-sdmm.dfo-mpo.gc.ca/isdm-gdsi/waves-vagues/data-donnees/data-donnees-eng.asp>) and buoy 45012 data from NOAA (<http://www.ndbc.noaa.gov/>) to calibrate and validate the integrated spectral wave (SW) module (Fig 1). This module was added to the modelling framework to determine the combined bottom shear stresses with those from the currents, but it also improved the prediction of surface currents in the nearshore area by including wave-generated currents due to breaking. The total bottom shear stresses were needed for calculating the resuspension of sediments and their associated nutrients in a related study about the role of *Dresissena* mussels and *Cladophora* in the nearshore nutrient cycle.

The tributary data was obtained from the United States Geological Survey (USGS) (<https://waterdata.usgs.gov/nwis/uv>) and for the Canadian lake side from the Canadian Hydrographic Service (https://wateroffice.ec.gc.ca/). Lake Ontario outflows were obtained from the International Lake Ontario-St. Lawrence River Board (<https://ijc.org/en/loslrb/watershed/outflow-changes>). Bathymetry data was obtained from the US National Geophysical Data Center (NGDC, 2009), which provides a 3 arc-second resolution representation of the lake (Fig 1). Supplementary higher resolution data, for Toronto Harbour, Humber Bay and Whitby Harbour areas, were obtained from local surveys done by the Toronto and Region Conservation Authority.

#### Part II. Calibration and Validation

We calibrated the Lake Ontario model using two sets of data for a period from January to December with our measured data (water temperature and currents) available from April to November and water levels and wave data available from third party sources for the entire year. This approach gave the thermal model the first three months of simulated time to warm up from the initial conditions of 4°C and still give the model enough data for calibration and validation. The calibration was done with data from year 2013 (instrument locations not shown) and the validation was performed with data from 2018 for the entire year. The calibration and validation accuracy were evaluated using visual inspection of long time series and metrics that included the Pearson correlation coefficient R and Normalized Root Mean Square Error, defined as:

$r_{OS}= \frac{N\sum_{1}^{N} O_{i}S_{i}-\sum_{1}^{N} O_{i}\sum_{1}^{N} S_{i}}{\sqrt{N\sum_{1}^{N} {O_{i}}^{2}-{(\sum_{1}^{N} O_{i})}^{2}} \sqrt{N\sum_{1}^{N} {S_{i}}^{2}-{(\sum_{1}^{N} S_{i})}^{2}}}$, Eq. 1

$NRMSE=\frac{RMSE}{\frac{1}{N}\sum_{1}^{N} S_{i}}$, where $RMSE=\sqrt{\frac{1}{N}\sum_{1}^{N} {{(O}_{i}-S_{i})}^{2}}$, Eq. 2

where *O_i_* are the observation sample points, *S_i_* are the simulated sample points indexed with *i,* and *N* is the sample size. The sample mean is $\bar{x}= \frac{1}{N}\sum_{1}^{N} x_{i}$ and the analog expression for $\bar{y}$. were used to simplify the resulting Eq 1.

For vector measures such as velocities, a normalized RMSE (F-NORM) for the components of the vectors was used:

$F_{N}= NRMSE =\frac{\sqrt{\sum_{1}^{N} {(({Xo}_{i}-{Xs}_{i})}^{2}+\left( {Yo}_{i}-{Ys}_{i} \right)^{2})}}{\sqrt{\sum_{1}^{N} \left( {{Xs}_{i}}^{2}+ {{Ys}_{i}}^{2} \right)}}$, Eq. 3

where *X_Oi_* and *Y_Oi_* are the eastward and northward measured velocity component sample points, indexed with *i*, and *X_Si_* and *Y_Si_* are the modelled eastward and northward velocity component sample points indexed with *i*, and *N* is the sample size.

Sensitivity analysis is of great importance to evaluate and quantify the uncertainties of models, including boundary conditions uncertainties, but also to identify the factors that have the most influence on model’s output, which in turn, minimize the calibration workload. Since in our model the boundary conditions were given, we determined the most influential parameters with short simulations by modifying one parameter at a time within an accepted range. We altered the parameters affecting the horizontal and vertical equations of motion, wave height, and those related to conservation of energy. A total of 19 parameters were considered (Table 1). Several meshes with different resolutions on the horizontal domain specification, and several vertical meshes configurations with Sigma layers only, and combinations of Sigma and Z-layers with different Sigma depths have been considered during sensitivity analysis and calibration phases. We tested combinations of high order and low order solutions for the shallow water equations, the thermal module, and the turbulence module.

**Table 1.** Range of parameters for the Lake Ontario model used for calibration after sensitivity analysis

| **Parameter** | | **Unit** | **Value of parameter**  *(min to max)* |
| --- | --- | --- | --- |
| Number of water layers | |  |  |
| Sigma layers only | |  | 20, 30 |
| Sigma depth | |  | 20, 40, 60 |
| Sigma + Z-layers | |  | 20S+10Z, 16S+14Z |
| Water friction parameters: | |  |  |
| Wind drag coefficient^1^ | | - | i) 0.0012 to 0.0026  ii) (see note below)  iii) 0.0024 to 0.0036 |
| Bottom roughness height | | m | *0.01 to 0.30 and variable* |
| Horizontal eddy viscosity: | |  |  |
| Smagorinsky coefficient, (*c_s_*) | | - | *0.10 to 1.0* |
| Horizontal eddy viscosity - upper limit | | m^2^s^-1^ | 100 to 1000 |
| - lower limit | | m^2^s^-1^ | 1.6 x 10^-6^ to 1.0 x 10^-5^ |
| Vertical eddy viscosity: used “Log law formulation” | |  |  |
| Upper limit | | m^2^s^-1^ | 0.1 to 2 |
| Lower limit | | m^2^s^-1^ | 1.8 x 10^-6^ to 1.8 x 10^-5^ |
| Minimum depth Cutoff | | m | -2.0 to -0.2 |
| Dalton's Law: | |  |  |
| constant *a1* | | ms^-1^ | 0.35 to 0.65 |
| wind coefficient *b1* | | - | 0.65 to 1.15 |
| critical wind speed | | ms^-1^ | 2 to 4 |
| Angstrom's Sun Constant Law: | |  |  |
| constant *a2* | | - | 0.1 to 0.4 |
| constant *b2* | | - | 0.35 to 0.85 |
| Beer's Law Equation: | |  |  |
| coefficient *λ* | | m^-1^ | 0.15 to 0.6 |
| coefficient *β* | | - | 0.15 to 0.6 |
| *Notes:* | | | |
| *(1)* | The wind drag coefficient *C_d_* is divided into 3-segments, according to wind speed (*U_w_*), as follows:  i) if: *U_w_* < 5 ms^-1^; then: *C_d_* = 0.0026  ii) if: 5 ms^-1^ < U_w_ < 15 ms^-1^ ; then: C_d_ = 0.0026 + 0.00010 * (U_w_ - 5)  iii) if: U_w_ > 15 m/s; then: C_d_ = 0.0036 | | |

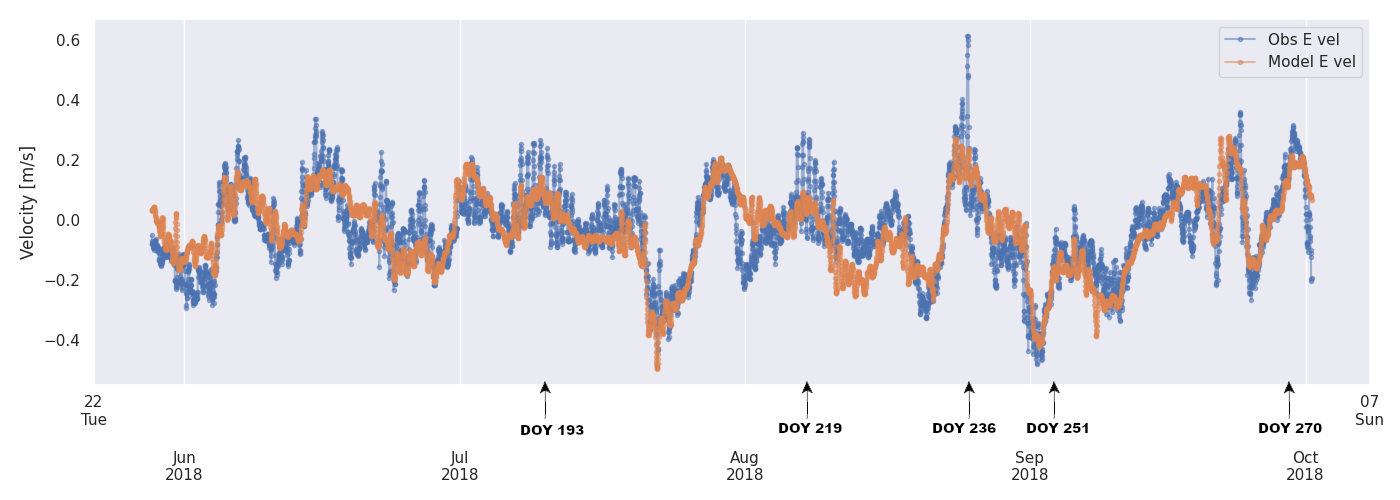

**S2 Fig.** Observed and Modeled E Velocity at Station E2_N20 at a depth of 6 m.

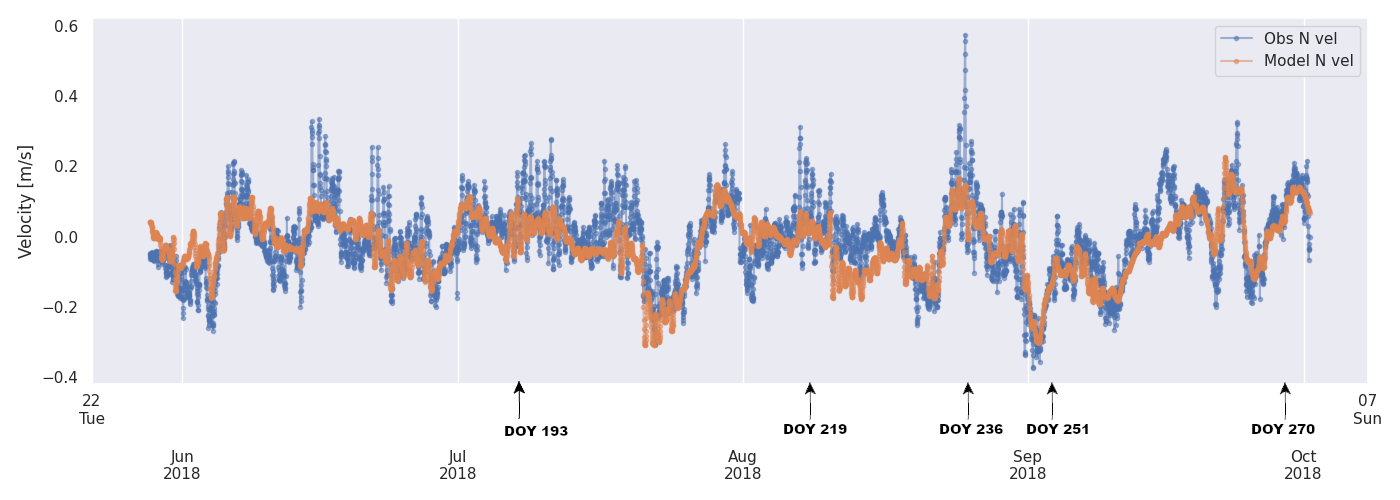

**S3** **Fig.** Observed and Modeled N Velocity at Station E2_N20 at a depth of 6 m.

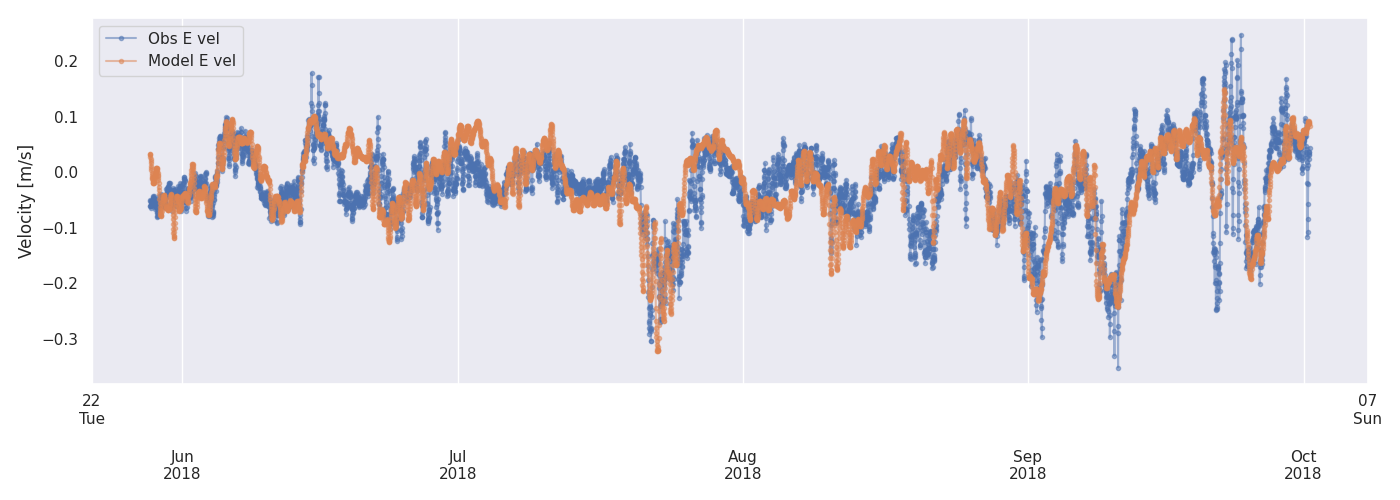

**S4** **Fig.** Observed and Modeled E Velocity at Station E2_N20 at a depth of 24 m.

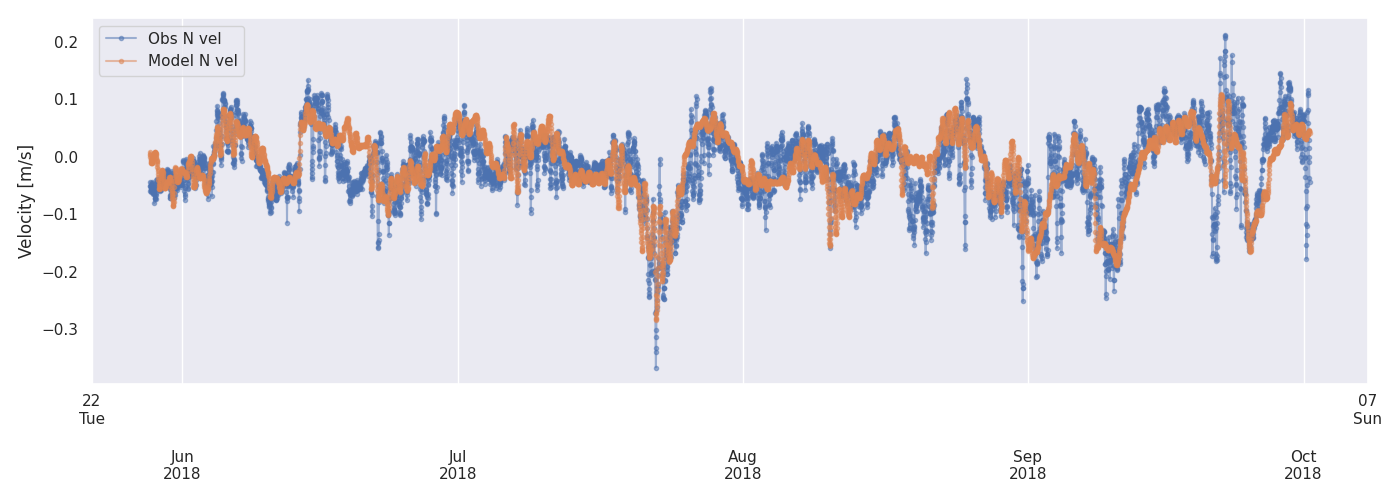

**S5 Fig.** Observed and Modeled E Velocity at Station E2_N20 at a depth of 28 m.

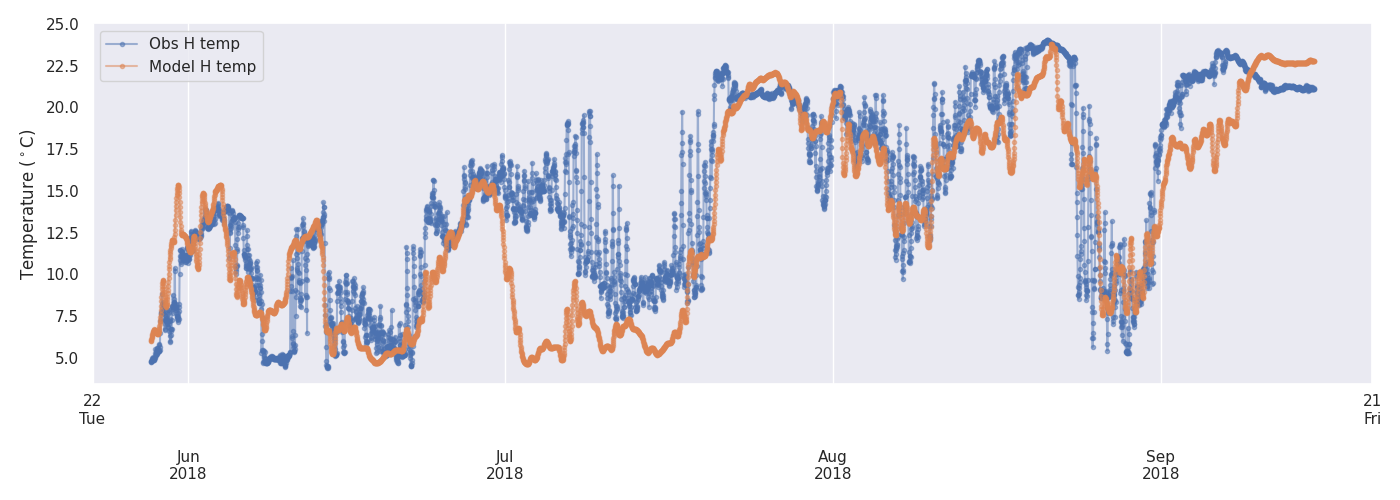

**S6** **Fig.** Observed and Modeled temperature at Station E2_N20 at a depth of 9 m

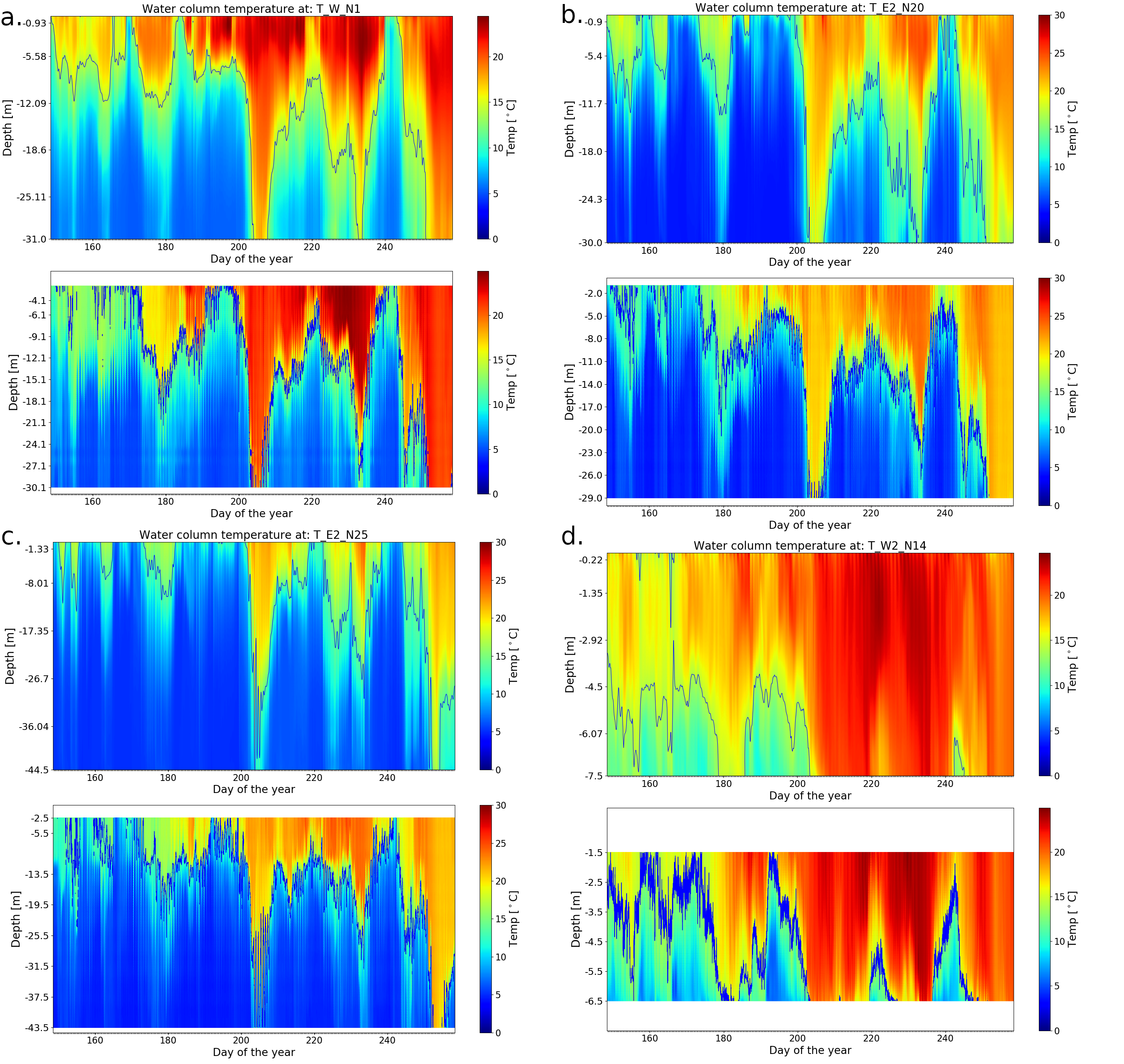

**S7 Fig**. Modeled temperatures and observations in 2018. The upper plot in each sub-figure is the model result and the bottom plot is the observed data. Data from the western station W2_N14 is incomplete due to gaps in the observations in the top layers. a. Station W_N1; b. Station E2_N20; c. Station E2_N25; d. Station W2_N14. The figures have different colour bar scales to optimize for colour representation.

### Part III. Additional data

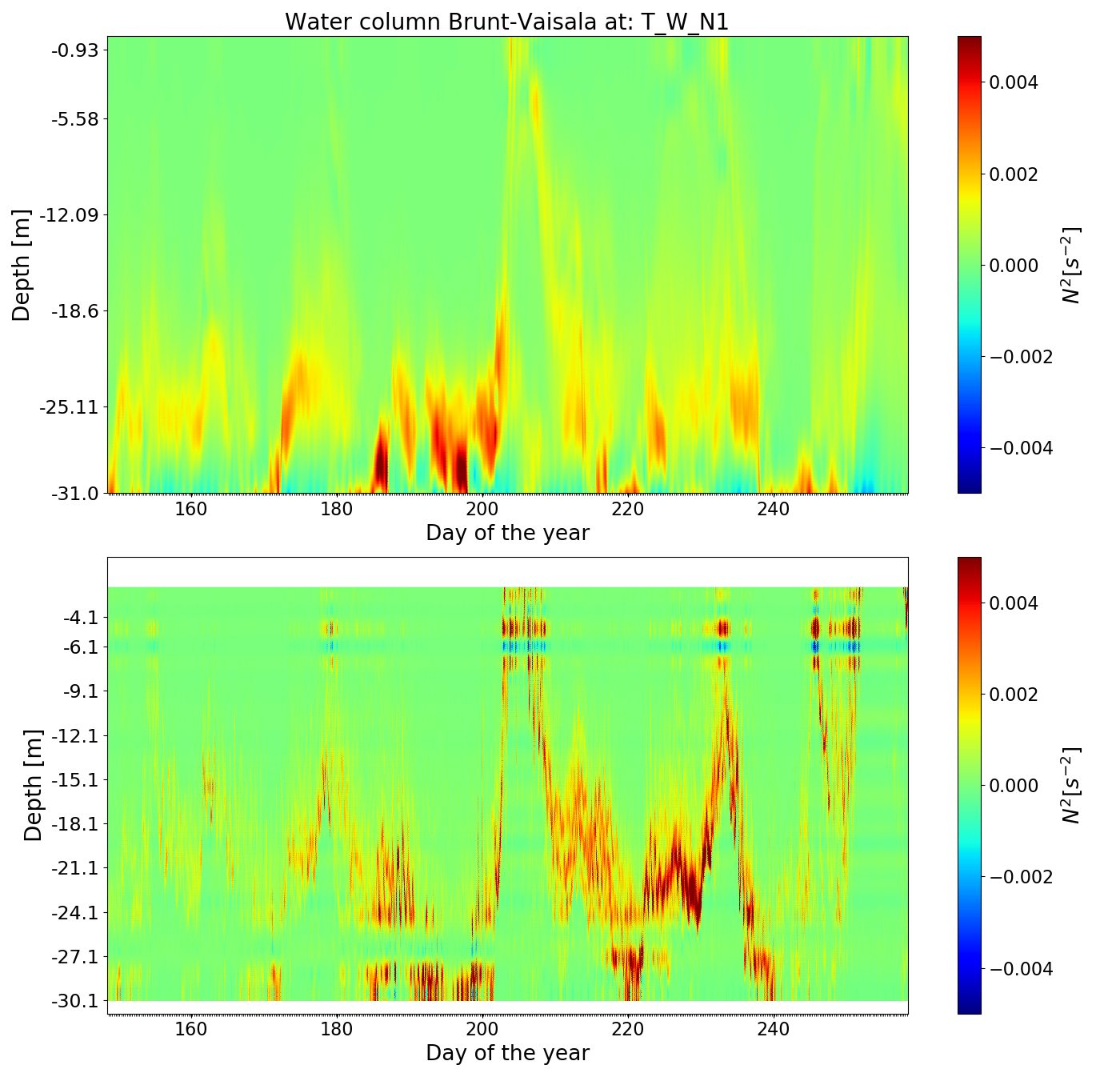

### S8 Fig. Brunt–Väisälä frequency in the water column at Station W_N1. a) calculated from the model result ; b) calculated from the observational data. The value of *N^2^* follows closely the position of the thermocline. The water column is stable (N2>0) during the statified season and starts crossing into the unstable zone after DOY 250

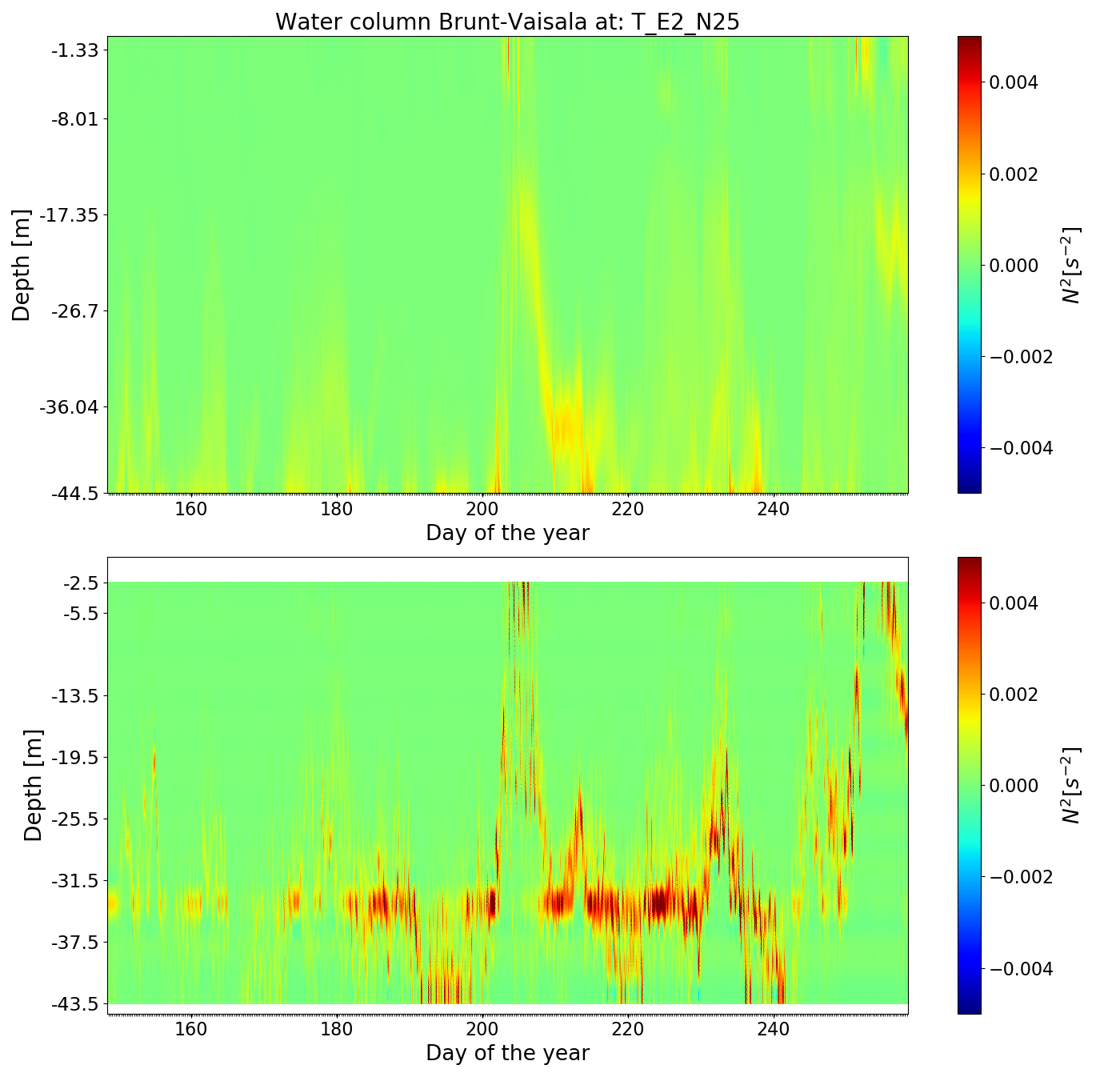

**S9 Fig**. Brunt–Väisälä frequency in the water column at Station E_N25. a) N^2^ estimated from observed data. b) N^2^ estimated from modelled data. The value of N^2^ follows closely the position of the thermocline.

### Part IV. Bottom slope analysis

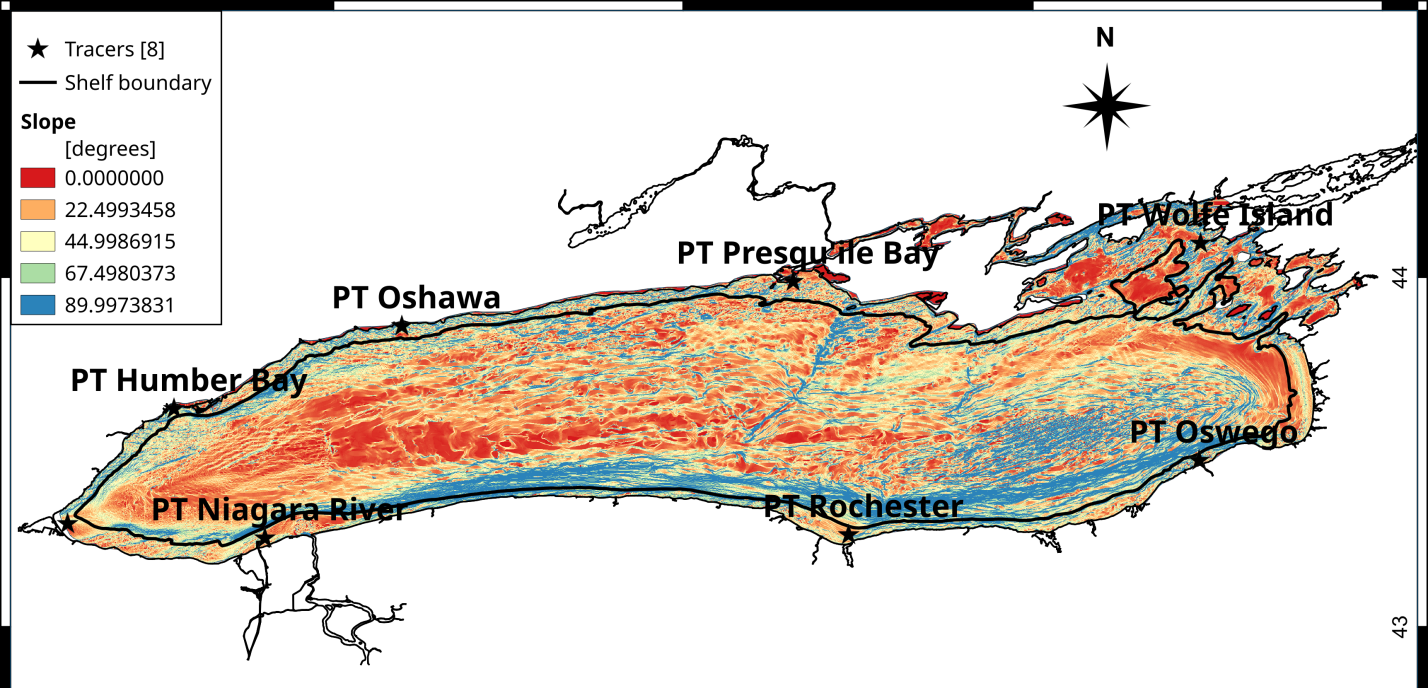

**S10 Fig.** Bottom slope map. The colours represent the slope values in degrees, from flat (red) to vertical (blue)

### Part IV. Field Observations

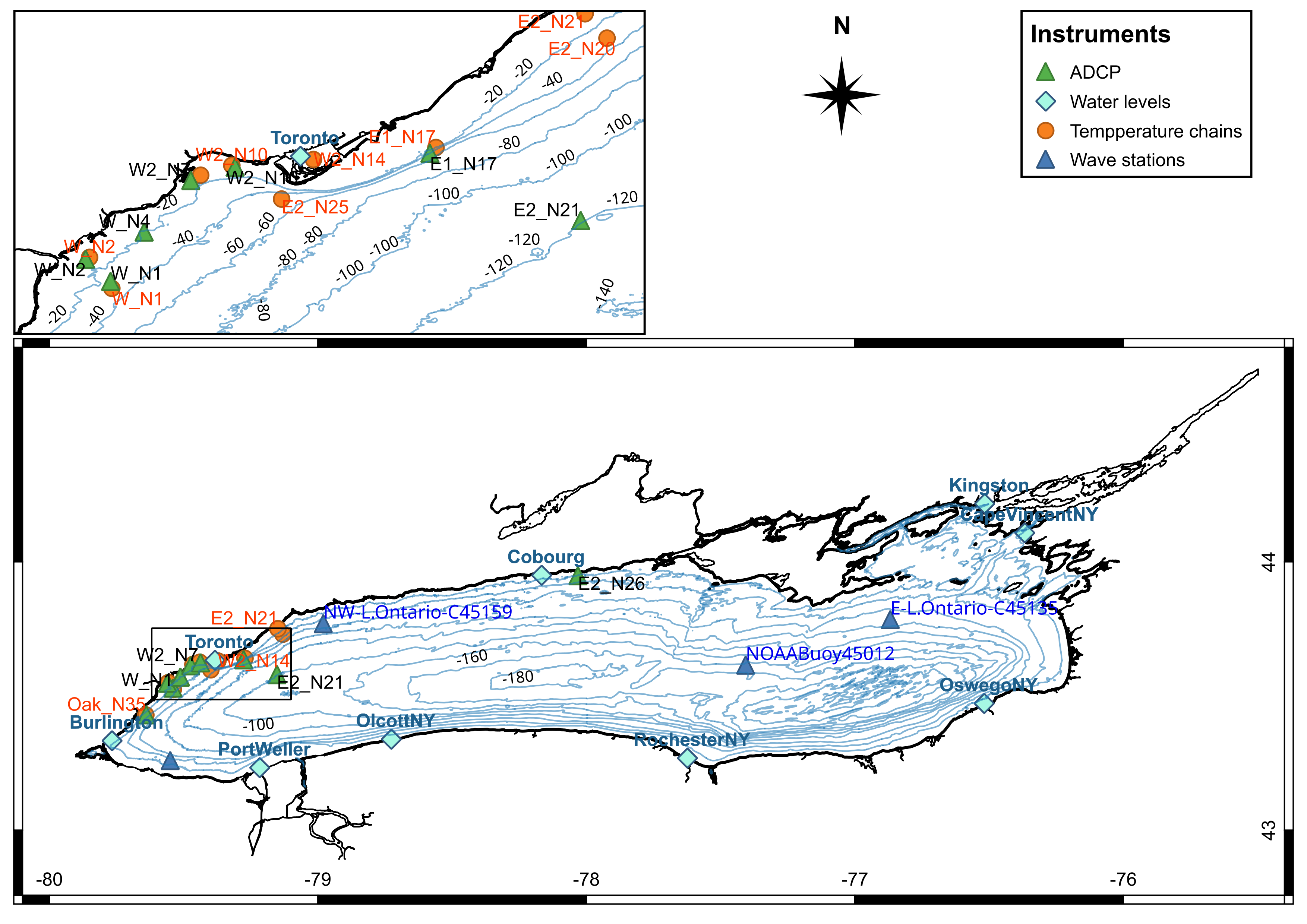

**S11 Fig**. Map of Lake Ontario, showing the locations of the deployments in the 2018 field campaign. The three offshore weather buoys are identified using blue triangles. Blue diamonds denote permanent water level gauges, green triangles stations with ADCPs, and orange dots denote locations with thermistor chains. The upper figure depicts the zoomed in overview of the Toronto waterfront, which hosts the majority of the stations.

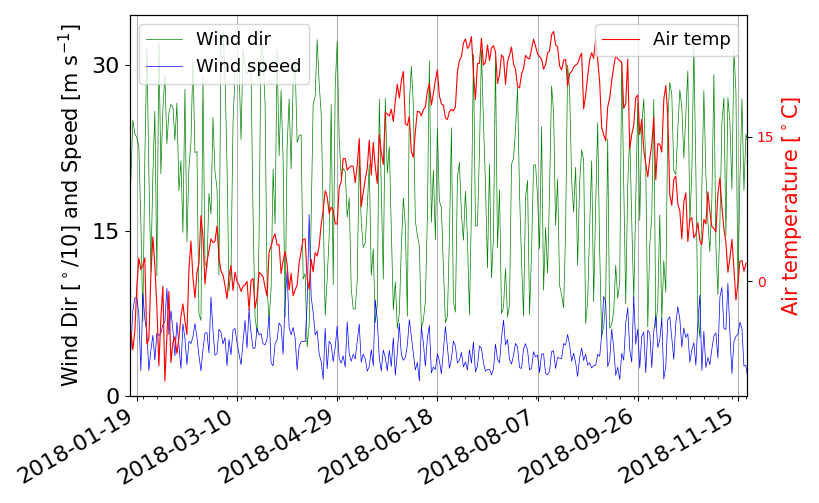

**S12 Fig** Wind speed and direction, and air temperature at Toronto Island Environment and Climate Change Canada station during the year 2018
